# *Clostridioides difficile* Toxins Alter Host Metabolic Pathway and Bile Acid Homeostasis Gene Expression In Colonic Epithelium

**DOI:** 10.1101/2025.03.21.644594

**Authors:** Stephanie A Thomas, Colleen M Pike, Cypress E Perkins, Sean T Brown, Xochilt M. Espinoza Jaen, Arthur S McMillan, Casey M Theriot

## Abstract

A major risk factor for acquiring *Clostridioides difficile* is antibiotic usage that disrupts a healthy microbial gut community facilitating establishment of infection. Once established, *C. difficile* secretes endotoxins (TcdA and TcdB) that are internalized into host colonic epithelial cells where they disrupt gut barrier function and induce hyper inflammation resulting in severe diarrhea and possibly leading to death. We employed three different platforms to explore gene expression of cells in the gut when exposed to *C. difficile* or its toxins, TcdA and TcdB. An antibiotic treated mouse model of *C. difficile* infection (CDI) was used to identify differential gene expression with a NanoString Technologies mouse inflammatory gene panel consisting of 770 genes including a subset of bile acid (BA) homeostasis and nuclear receptor genes. In the cecal tissue of mice with CDI, significant down expression was observed for genes involved in PPAR signaling, cholesterol and glucose metabolism, while a significant increase in expression was observed for IL-17 related inflammatory genes. Similarly, Caco-2 cell culture and primary human colonocytes (hCE) exposed to toxins for 24 hours showed altered expression in several PPAR regulated and cholesterol metabolic genes similar to those found in mice. These cell culture experiments also revealed significant alterations in gene expression of the FXR BA regulatory pathway. Together these data suggest that exposure to *C. difficile* and its toxins may alter host cholesterol metabolic processes, including BA transport and synthesis.

## Introduction

*Clostridioides difficile* infection (CDI) remains an urgent public health threat by the CDC with an incidence of approximately 500,000 cases per year in the US and 29,000 deaths^1-3^. A major risk factor for acquiring *C. difficile* is antibiotic usage that disrupts a healthy microbial gut community facilitating CDI establishment^4,5^. Once colonization is established, *C. difficile* secretes toxins (TcdA and TcdB) that are internalized in colonic epithelial cells where they inactivate Rho and Rac GTPases by glucosylation^6-8^. Inactivation of these small GTPases disrupts the actin cytoskeleton and cellular tight junctions, ultimately causing apoptosis or necrosis^6^. Immune cell infiltration is important in defense against CDI, however, the prolonged influx of pro-inflammatory mediators leads to increased tissue damage that can result in severe diarrhea, toxic megacolon, pseudomembranous colitis and even death^6,9^. Current treatments include subsequent antibiotic treatments such as vancomycin and fidaxomicin, leaving limited treatment options for the 30% of patients that develop recurrent CDI (rCDI). For some patients, introducing intact, healthy microbiota from a fecal microbiota transplantation (FMT) and or using newer FDA approved microbiota focused therapeutics like Vowst and Rebyota are their last resort.

*C. difficile* is exquisitely sensitive to gut specific bile acids (BAs). Spore germination occurs in the presence of taurocholate (TCA), a primary BA produced by the liver^10^. Alternatively, *C. difficile* growth is sensitive to secondary BAs (primary BAs modified by gut microbes), namely deoxycholate (DCA) and lithocholate (LCA)^11,12^. The loss of gut bacteria due to antibiotic usage depletes secondary BAs and is associated with colonization of *C. difficile*. Recovery of secondary BAs in the intestinal BA pool is observed with successful FMT treatment^13,14^. The primary regulator of BA synthesis is the nuclear receptor Farnesoid X Receptor (FXR). FXR is activated by high levels of BAs in the intestinal lumen. Activation of FXR leads to increased expression of key players in BA homeostasis, including BA transporters (FABP6, ASBT, OSTα/β) and FGF19, the secreted signaling molecule that triggers down regulation of BA production in the liver creating a negative feedback loop^15^. FXR activation also protects against cell damage by stabilizing barrier integrity and reducing inflammation by inhibiting transcription of NF-kB regulated cytokines such as IL-1β^16-19^. Several BAs serve as FXR agonists including the primary BA chenodeoxycholate (CDCA) and secondary BAs, DCA and LCA and their conjugated derivatives^20^. Enriching the intestinal BA pool with BAs that function as FXR agonists may enhance FXR activity, reduce toxin-mediated inflammation in CDI, and serve as a promising therapeutic strategy during infection.

It is well established that *C. difficile* toxins induce changes to the host cellular structure and inflammatory response^6-8^, but little is known about BA modification during CDI. Previous work has shown that *C. difficile* triggers an influx of TCA that accumulates in the ileal lumen, altering the BA pool in a toxin dependent manner^21^. The underlying mechanisms of this finding remain unclear. A deeper understanding of the interaction between *C. difficile* and BA regulation is needed before exploring FXR agonists as a potential therapeutic. In this study we leveraged two *in vitro* and one *in vivo* platform to assess gene expression alterations in host BA homeostasis genes when exposed to *C. difficile* or purified toxins. Platforms included a primary CDI mouse model, toxin exposure to Caco-2 cells, and primary colonocytes derived from the intestinal crypts of a healthy human donor (hCE). Differential gene expression (DE) in genes of interest including FXR and its target genes involved in BA homeostasis and cholesterol transporters were evaluated. Our findings show mice challenged with *C. difficile* had reduced gene expression in metabolic pathways, including glycolysis, cholesterol metabolism, and PPAR signaling, and increases in several inflammatory genes in the gut. Gene expression of select genes identified in mice were also decreased in the hCE cells, but increased in the Caco-2 immortal cell line when exposed to toxins alone. *C. difficile* toxin exposure can alter expression of BA homeostasis genes without FXR activation and suggests that toxins could be modifying BA and cholesterol transport in colonocytes.

## Methods

### Animals and housing

C57BL/6J mice (males) were purchased from Jackson Laboratories (Bar Harbor, ME) and quarantined for 1 week prior to starting the antibiotic water administration to adapt to the new facilities and avoid stress-associated responses. Following quarantine, the mice were housed with autoclaved food, bedding, and water. Cage changes were performed weekly in a laminar flow hood by laboratory staff. Mice had a 12-hr light and dark cycles, and were housed at an average temperature of 70°F and 35% humidity. NCSU CVM is accredited by the Association for the Assessment and Accreditation of Laboratory Animal Care International (AAALAC). This protocol is approved by NC State’s Institutional Animal Care and Use Committee (IACUC).

### Mouse model of *C. difficile* infection and sample collection

Methodologies of this primary CDI model have been previously published^22,23^. Groups of 5-week-old C57BL/6J mice (male; 4 mice/treatment group) were given cefoperazone (0.5 mg/ml) in their drinking water ad libitum for 5 days, followed by a 2-day wash out with regular drinking water ad libitum (Fig. 1A). Mice were then challenged with 10^5^ spores of *C. difficile* R20291 pyrE via oral gavage at day 0. Mice were monitored for weight loss and clinical signs of CDI (lethargy, inappetence, diarrhea, and hunched posture) for 2 days post challenge. Fecal pellets were collected daily for *C. difficile* enumeration. Uninfected controls included cefoperazone-treated mice and conventional mice. Animals were euthanized 2 days post challenge. The contents and tissue from the cecum and colon were collected immediately at necropsy, flash frozen in liquid nitrogen, and stored at -80°C until further analysis. To prevent RNA degradation, snips of cecal and colonic tissue were also stored in RNA-Later at -80°C until further analysis.

**Figure 1.**
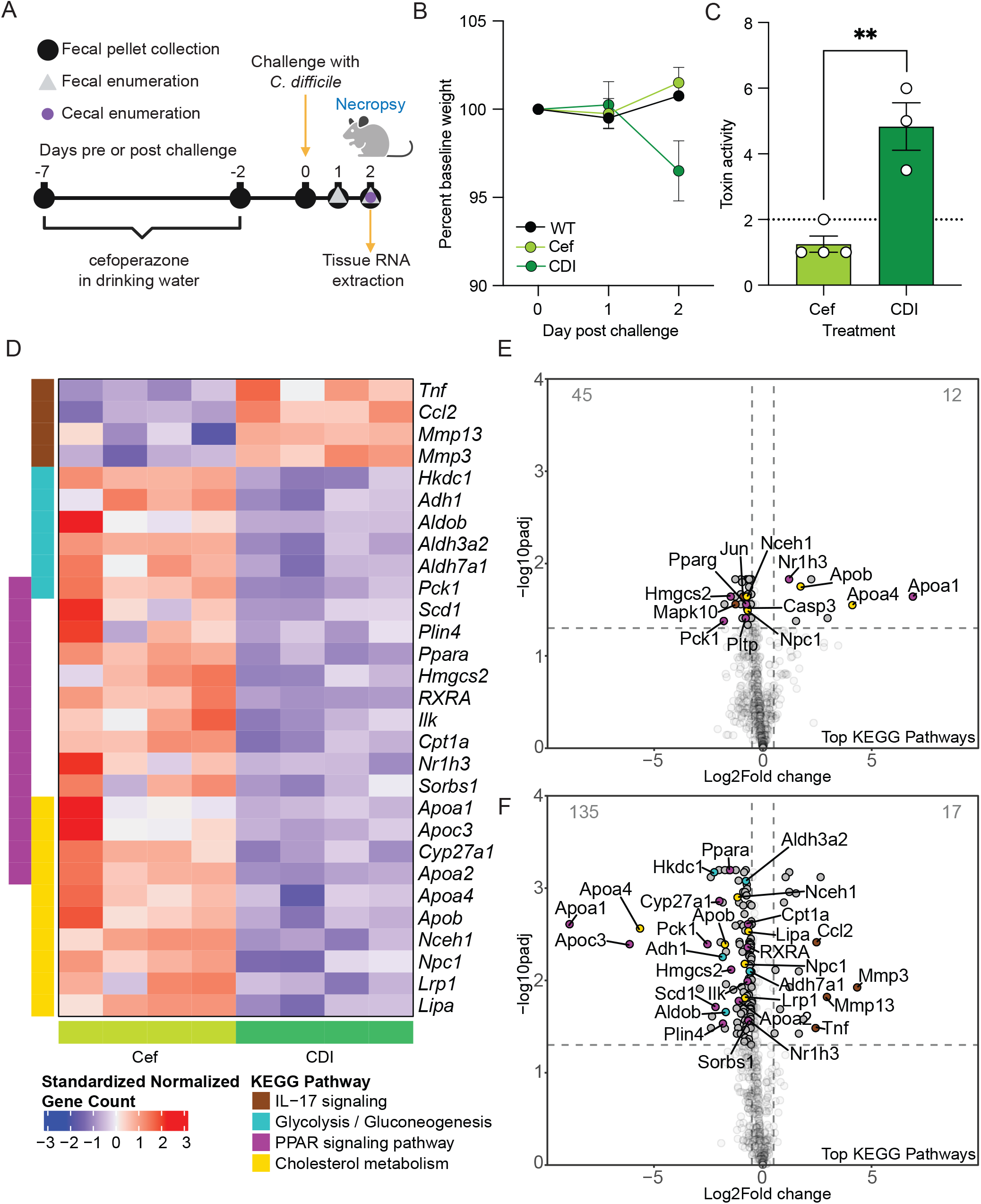
Differential gene expression in mouse cecal tissue during *C. difficile* infection compared to antibiotic treated mice. A) Mouse model of CDI experimental details. Mice (n=4 per treatment group) were exposed to one of three treatments, no treatment or wild type (WT), cefoperazone treatment only (Cef), and cefoperazone treated mice followed by challenge with 10^5^ spores of *C. difficile* (CDI). B) Percent baseline weight loss of mice from day 0-2 post challenge. C) Toxin activity assay from the cecal content of CDI mice and Cef mice. An unpaired t-test was used for statistical analysis with \*\**p* < 0.01 (*p* = 0.0032). D) Significantly DE genes clustering to one of the 4 KEGG pathways identified in the gene set enrichment analysis are represented in heatmap as gene counts from individual mice treated with Cef (light green) or CDI (dark green). E) Volcano plots of genes differentially expressed in Cef mice compared to WT mice or F) CDI mice compared to Cef mice. Colored circles correspond to the KEGG pathways represented in D. Differential expression analysis was performed using nSolver Advanced analysis software v2.0.134 using Benjamini-Hochberg p-value adjustment with *p*<0.05.

### Vero cell cytotoxicity assay

Toxin activity was tested as described previously^22^. Cecal contents were resuspended in 1:10 dilution w:v in PBS and filtered with a 0.22 μm filter. Filtrates were added to wells seeded with 10^4^ cells per well and incubated overnight. Toxin A from VPI 10463 was used as a positive control and antitoxin (List Labs 152C, and TechLab, T5003) was used to quench activity.

### RNA isolation and gene expression analysis

RNA extraction was performed on cecal tissue using PureLink RNA Mini kit (Thermo Fisher, 12183025) following the manufacturer’s protocol. The RNA was treated with Turbo DNase (Thermo Fisher, AM2239) to remove genomic DNA contamination. Samples were quantified using Qubit (Thermo Fisher Scientific) and run on an Agilent Bioanalyzer NanoChip (Santa Clara, CA) to assess the quality of RNA. The nCounter Mouse Fibrosis v2 gene expression panel and custom code set was purchased from NanoString Technologies (Seattle, WA) and consists of 770 mouse genes, including six internal reference genes and 19 custom genes of interest (Table S1). The nCounter assay was performed using 100 ng of total RNA. Hybridization reactions were performed according to the manufacturer’s instructions with 5 μl diluted sample preparation reaction, and samples were hybridized overnight. Hybridized reactions were purified using the nCounter Prep Station (NanoString Technologies), and data collection was performed on the nCounter Digital Analyzer (NanoString Technologies) following the manufacturer’s instructions to count targets. Nanostring nCounter Max platform was utilized at the Microbiome Shared Resource Lab at Duke University to perform these assays.

Raw data from NanoStrings nCounter was analyzed using the nSolver Analysis Software v4.0. Gene counts for each sample were background subtracted based on the geometric mean of negative probes. Gene counts for each sample were normalized based on the geometric mean of positive probes and to a panel of housekeeping genes; *Acad9, Armh3, Cnot10, Gusb, Mtmr14, Nol7, Nubp1, Pgk1, Ppia, Rplp0*. Differential expression analysis was performed using nSolver Advanced analysis software v2.0.134 using Benjamini-Hochberg p-value adjustment. Probes used custom annotations as detailed in Table S2. Further analysis was performed in R v4.4.1. Ensembl and NCBI gene IDs were annotated using biomaRt v2.60.1 with the GRCm39 gene ensemble. Gene set enrichment analysis (GSEA) was performed on differentially expressed genes using a minimum gene set size of 3 using clusterProfiler v2.1.6 to perform GSEA with Benjamini-Hochberg correction. Graphing was performed using tidyverse v2.0.0 and ComplexHeatmap v2.20.0. Normalized gene counts in heatmaps were standardized by subtracting the mean and dividing by the standard deviation for each probe. All computational analysis is available doi: <u>10.5281/zenodo.13350814</u>.

### RNA extraction from Caco-2 cells

24 well plates were seeded with 5x10^4^ Caco-2 (ATCC, HTB-37) cells per well and grown in DMEM supplemented with 2 mm L-glutamine, 10% FBS, and penicillin 5000 U/ml plus 5000 μg/ml streptomycin cocktail. Plates were incubated in 5% CO_2_ at 37 °C with media changes every 2-3 days. After cells reached confluency (3-4 days) they were incubated for an additional 7 days. After 7 days post confluency, cells were treated with 100 pmol of recombinantly purified TcdA, TcdB, or TcdB lacking the Gycosyl-transferase domain activity (TcdB-GTD^mut^)for 24 hr before RNA extraction. TcdB-GTD^mut^ mutations in the GTD domain at the following residues, W102A/D286N/D288N. This mutant also lacks residues 2101-2366 in the C-terminus of the CROP domain allowing complete exposure of the receptor binding domain (DRBD). After incubation, cells were lysed with Trizol reagent. 200 μl of chloroform was added to the lysate mixed and incubated for 15 min. The lysate was centrifuged at 4° C for 15 min and the aqueous layer was removed added to equal volumes of 100% ethanol. The RNA from the ethanol mixture was purified using the Zymo Research RNA Clean and Concentrator-25 (Zymo Research, Cat# R1017) per manufacturer’s instructions. DNA was degraded on the column with Zymo Research DNAse 1 (Zymo Researach, Cat# E1010) per manufacturer’s protocol. RNA concentration was measured with Qubit.

### RNA extraction from human colonocytes

Primary human transverse colon epithelial cells were acquired from Altis Biosystems and cultured as previously described^24,25^. Cells were plated onto a 96-well Transwell® plate (Corning, 3392) and grown in Altis RepliGut® Growth Media (RGM). Once cells reached confluence, Altis RepliGut® Maturation Medium (RMM) was applied to promote cellular differentiation and polarization. hCE barrier integrity was monitored daily via TEER using the EVOM™ Auto (EVA-MT-03-01, WPI). Treatments used for this study were 1) no treatment-cells were grown in RMM only, 2) vehicle-RMM media with 2% vehicle buffer (20 mM Tris pH 7.5, 150 mM NaCl), 3) obeticholic acid (OCA) added to RMM at a concentration of 200 nM, 4-5) TcdA or TcdB - toxin diluted in vehicle and added to RMM to a final concentration of 100 pM toxin and 2% vehicle buffer. At 24 hr post-treatment, RNA lysates were collected in Buffer RLT (Qiagen, Cat# 79216). RNA extraction followed the Qiagen Rneasy Plus extraction kit (Qiagen, Cat# 74134).

### Quantitative reverse transcription PCR

RNA from Caco-2 cells and human colonocytes were normalized to 500 ng and 100 ng respectively and used as a template for reverse transcription reactions using the High-Capacity cDNA Reverse Transcription Kit (ThermoFisher Cat# 4368814). The resulting cDNA was used for quantitative PCR with the SsoAdvanced Universal SYBR Green Supermix (Bio-Rad Cat# 1725270). For relative quantification, the ΔΔCt method was used to normalize the gene of interest to the internal expression control, *GAPDH*, followed by normalization of toxin treated cells to no treatment controls. Data is reflected as a fold change comparison between toxin treated cells to no treatment controls.

### Protein expression and purification of TcdA and TcdB

Expression and isolation of recombinant TcdA, TcdB and TcdB-GTD^mut^ was performed as described by Yang et al.^26,27^. Purified proteins were a gift from Roman Melnyk and John Tam. Briefly, transformed *Bacillus megaterium* was inoculated into LB containing tetracycline and grown to an A_600_ of 1.6, followed by overnight xylose induction at 30°C. Bacterial pellets were collected, resuspended with 20 mM Tris pH 8/0.1 M NaCl, and passed twice through an EmulsiFlex C3 microfluidizer (Avestin, Ottawa, ON) at 15,000 psi. The resulting lysate was clarified by centrifuging at 18,000 ×g for 20 min. TcdB and TcdA were purified by nickel affinity chromatography followed by anion exchange chromatography using HisTrap FF Crude and HiTrap Q columns (Cytiva), respectively. TcdB-GTD^mut^ was purified similarly with the exceptions of initial purification used a cobalt affinity column and the eluent was biotinylated at 4°C with 15X molar excess maleimide biotin (Thermo 21901BID) and then followed by a HiTrap Q column. Fractions containing purified protein were verified by SDS-PAGE, then pooled and diafiltered with a 100,000 MWCO ultrafiltration device (Corning) into 20 mM Tris PH 7.5/150 mM NaCl. Finally, glycerol was added to 5% v/v, the protein concentration was estimated by A280, divided into single use aliquots, and stored at -80°C.

### Statistical analysis

An unpaired t-test was used for the toxin assay (Fig. 1C) statistical analysis (*p* < 0.05). For the toxin assay (Fig. 1C) and qRT-PCR data (Fig. 2A and B), statistical tests were performed in GraphPad Prism 8. For qRT-PCR data, normalization assays were conducted using the Shapiro-Wilk test. Differential expression analysis for NanoStrings data was performed using nSolver Advanced analysis software v2.0.134 using Benjamini-Hochberg p-value adjustment. Genes with a log2 fold change >0.5 or <-0.5 and *p* < 0.05 were called as differentially expressed genes (Fig. 1E and F). Caco-2 expression data was not normally distributed (Shapiro-Wilk test) and treatments were compared to no treatment (NT) control using the Mann-Whitney non-parametric test (Fig. 2A and Suppl. Fig. 2). hCE data showed normal distribution and an unpaired t-test was used to compare treatments to NT (Fig. 2B). Significance for qRT-PCR analysis is *p* < 0.05.

**Figure 2.**
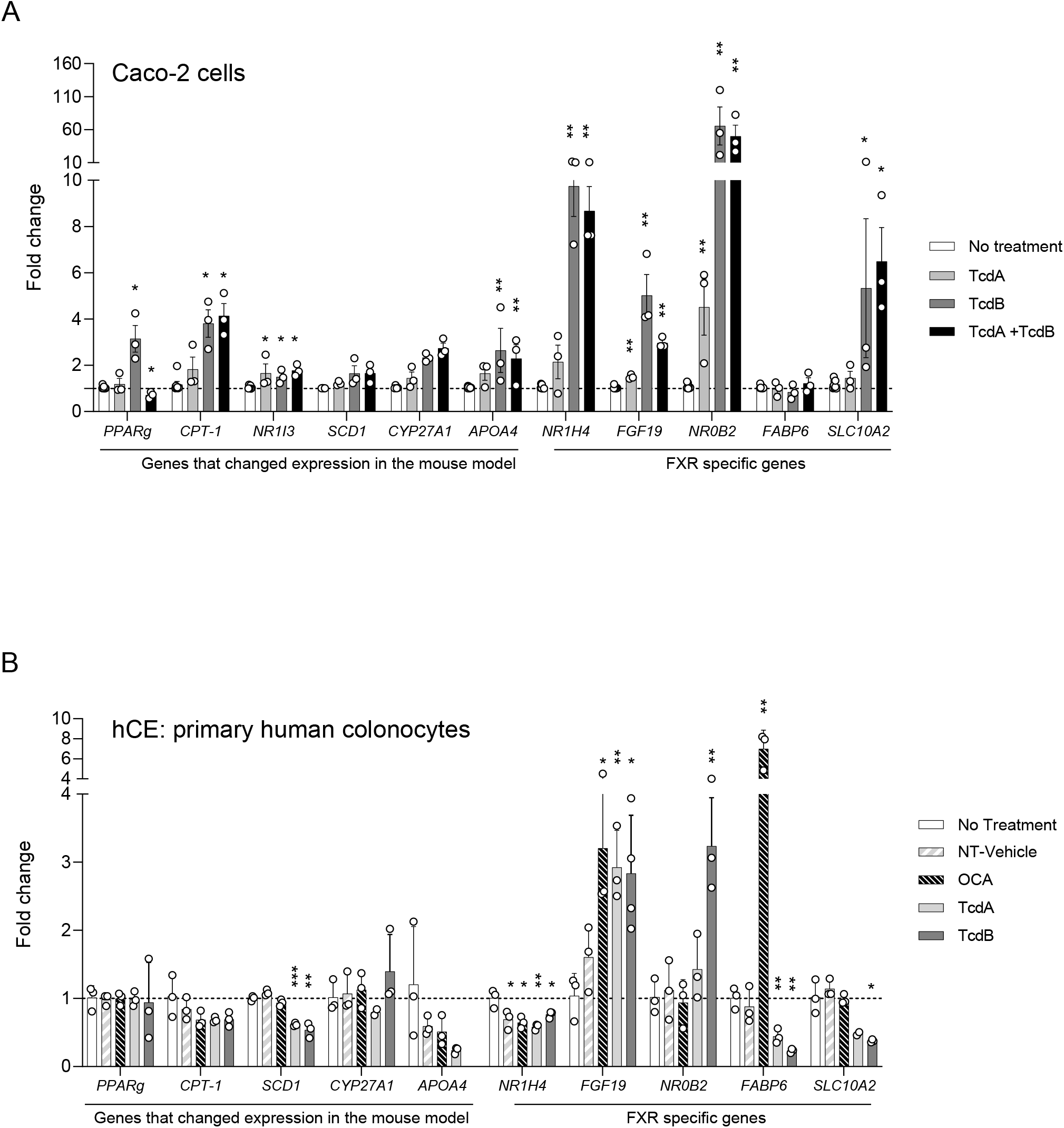
Toxin exposure alters expression of PPAR and FXR signaling pathways in Caco-2 monolayers and primary human colonocytes. A) Caco-2 gene expression, wells were treated with 100 pM of either TcdA, TcdB, or 100 pM of both TcdA and TcdB and incubated for 24 hr. B) hCE gene expression, cells were treated with 100 pM of either TcdA or TcdB. 200 nM of OCA was also added as an FXR agonist and positive control. No treatment control consisted only of media with no other additions. 20 mM Tris pH 7.5, 150 mM NaCl diluted to 2% in RMM acted as the no treatment vehicle control. For A, asterisks denote statistical significance determined by Mann-Whitney t-test compared to the No treatment control. \**p* < 0.05, \*\**p* < 0.01. For B, asterisks denote statistical significance determined by unpaired t-test compared to No treatment control. \**p* < 0.05, \*\**p* < 0.01.

## Results

### CDI decreases gene expression of host cholesterol metabolism in mice

An antibiotic treated mouse model of CDI was used to investigate changes in host gene expression during acute infection (Fig. 1A). CDI mice were symptomatic 2 days post challenge with *C. difficile* spores evidenced by weight loss compared to either wildtype (WT) or antibiotic treated mice (Cef) (Fig. 1B). Significantly greater toxin activity was observed in the cecal content of CDI mice compared to the Cef mice as expected^22^, confirming active CDI (Fig. 1C, *p*= 0.0032, unpaired t-test). CDI mice had a *C. difficile* cecal load of approximately 1.15x10^8^ CFU/g . At the time of necropsy, cecal tissue was collected from each treatment group for later RNA extraction and differential gene expression leveraging NanoStrings Technologies. We used the mouse inflammatory gene panel consisting of 770 genes and a subset of BA homeostasis and nuclear receptor genes. Gene set enrichment analysis (GSEA) was used to identify KEGG pathways with significant differentially expressed genes (DE) (Fig. S1). Four pathways were identified; the IL-17 signaling pathway consisted of four genes with increased expression (*Tnf, Ccl2, Mmp13*, and *Mmp3*) during CDI compared to Cef, while 25 genes with reduced expression were spread between glycolysis/gluconeogenesis metabolism, cholesterol metabolism, and PPAR signaling pathway (Fig. 1D, S1A-D). Most genes with reduced expression were associated with cholesterol metabolism where several genes are also regulated through PPAR signaling pathways (Fig. 1D, S1C-D). Volcano plots comparing host gene expression of Cef mice compared to WT mice (Fig. 1E) show a small number of genes with altered expression (12 increased, and 45 decreased), indicating that exposure to the antibiotic alone had a subtle effect on gene expression. However, volcano plots comparing host gene expression in CDI compared to Cef mice (Fig. 1F) indicate that exposure to CDI does significantly impact gene expression, with 17 genes significantly increased in expression and 135 significantly decreased in expression (Fig. 1F). Reduction in cholesterol transport genes were prominent, namely the apolipoproteins, *Apoa1, Apoa4, Apoc3* and *Apob* (Fig. 1D, 1F). BA related genes, *Cyp27a1* and *Scl51a* (OSTa) had significantly reduced expression in mice with CDI (Fig. 1D, 1F, Table S1). Expression of several nuclear receptors including VDR (*Nr1i1*), CAR (*Nr1i3*), LXRβ (*Nr1h2*), LXRα (*Nr1h3*), and *RXRA* (Table S1) was also significantly decreased in mice with CDI.

### Toxin exposure in Caco-2 cells and human primary colonocytes disrupts cholesterol and bile acid transport gene expression

To determine if toxins alone can modify transcription of the PPAR signaling pathway and cholesterol metabolism genes in human colon cells in a similar manner as mouse cecal tissue, we exposed differentiated Caco-2 cells grown in a flatbottom well system to toxins TcdA and TcdB or both at 100 pM for 24 hours. Caco-2 cells exposed to TcdA had significantly altered expression of the gene encoding CAR (*NR1I3*) (Fig. 2A). Caco-2 cells exposed to TcdB had significantly increased expression of PPAR signaling genes *PPARG*, and *CPT-1*, as well as the apolipoprotein *APOA4*, and the nuclear receptor CAR (*NR1I3*) (Fig. 2A). When both toxins (TcdA + TcdB) were combined at 100 pM each, similar increases in expression to TcdB only exposure were observed, indicating that TcdB is responsible for the changes in expression. This was not true for *PPARG*, where expression decreased upon exposure to both toxins (Fig. 2A).

*Cyp27a1* and *Slc51a* were the only BA homeostasis genes with significant changes in gene expression levels in the mouse study. Mice typically produce high levels of the FXR antagonist TβMCA^28^, potentially inhibiting FXR activation and in turn, inhibiting expression of the FXR regulated BA genes. We wanted to explore these BA homeostasis genes further in an antagonist free environment and used qRT-PCR to determine if toxins altered gene expression in Caco-2 cells. Significant increases in gene expression were observed in cells exposed to all three treatments, TcdA, TcdB, and TcdA + TcdB (Fig. 2A). TcdB and TcdA + TcdB exposed cells had significantly increased expression in *NR1H4* (FXR), *FGF19, NR0B2* (SHP), and *SLC10A2* (ASBT) genes, but not *FABP6*. TcdA exposed cells had significantly increased expression in *FGF19* and *NR0B2* only. *NR0B2* had the highest increase in expression when exposed to TcdB and TcdA + TcdB, with a 65-fold and 50-fold increase, respectively. All other genes tested had less than 10-fold increases in expression.

To determine if disruption of Rho-GTPase signaling is responsible for the observed changes in expression seen in only the FXR regulated genes we exposed Caco-2 cells to 100 pM of the TcdB mutant lacking glycosyl-transferase activity (TcdB-GTP^mut^), and yet maintains all other functional characteristics of the protein. DE seen with fully active TcdB is lost with the mutant protein (Supp. Fig. 2), indicating that TcdB is altering gene expression via disruption of GTPase signaling.

Caco-2 cells are derived from an immortal adenocarcinoma in which expression of *NR1H4* and *FABP6* differ from primary colonic epithelial cells^29,30^. To avoid confounding variables using cancer cell lines, we implemented the RepliGut® platform (Altis Biosystems, Durham, NC) comprised of differentiated primary colonic epithelial (hCE) cells derived from intestinal crypts of human transplant-grade donors^24,25^. hCE were exposed to differing treatments for 24 hours and expression levels for the same PPAR pathway and cholesterol genes assayed in the Caco-2 cells were measured. *SCD1* was the only gene to result in significant differences, however, instead of increasing in expression like the Caco-2 cells, gene expression decreased in hCE cells (Fig. 2B). In fact, expression of other PPAR pathway genes resulted in little change or decreased expression in the hCE cells (Fig. 2B).

Significant differences in gene expression were observed in the BA regulatory genes in hCE (Fig. 2B). *NR1H4* expression was significantly reduced when exposed to either TcdA or TcdB compared to the no treatment control. These data are opposite of *NR1H4* expression in Caco-2 cells. However, like the Caco-2 cells, *FGF19* expression was significantly increased in the hCE cells when exposed to either TcdA or TcdB alone, but only TcdB significantly increased *NR0B2* expression. Significant decreases in expression of *FABP6* were observed for both toxins, but only TcdB exposure resulted in significantly reduced *SLC10A2* expression. The addition of OCA, an agonist of FXR, at 200 nM did not increase *NR1H4* expression, but OCA exposure increased expression of *FGF19* and *FABP6*, both of which are induced by activated FXR, suggesting OCA exposure did activate FXR leading to increased expression of these genes. Interestingly, activation of FXR via OCA is expected to increase *NR0B2* expression, however OCA exposure resulted in no change. The DE observed for *NR0B2* and *FABP6* when exposed to toxin is not through FXR and suggests that TcdA and TcdB can alter expression of BA homeostasis genes without activation of FXR by an agonist.

## Discussion

It is well known that *C. difficile* is exquisitely sensitive to BAs, namely, TCA which acts as a spore germinant^31^. Wexler et al. has shown that TCA concentration increases in the intestinal lumen in the early stages of CDI and is dependent on toxin^21^. However, the mechanism by which *C. difficile* is able to modify the host response with regards to BA synthesis is not known. This study used three different systems to interrogate the host BA response to *C. difficile* toxin exposure. In this study we provide evidence that *C. difficile* is able to modify cholesterol transport and metabolism including BA transport in a toxin dependent manner.

Differential gene expression in the mouse cecum did not result in significant changes in BA regulatory genes. However, several genes that decreased in expression are important in cholesterol and lipid transport. These include apolipoprotein genes *Apoa1, Apoa4, Apoc3* and *Apob* (Fig. 1D, 1F). These apolipoproteins are important components of chylomicron and high-density lipoproteins (HDL) structure which pack and transport cholesterol and lipids to the lymph system from intestinal epithelial cells^32-34^. The reduction in gene expression of intestinal apolipoproteins hints at reduced lipid and cholesterol transport into the lymph and blood stream. *Ppara* was also significantly reduced in cecal tissue and correlates with the reduced apolipoproteins as PPARs are transcriptional regulators of *Apoa1, Apoc3, Apoa2*^35,36^. Further support for disruption of lipid transport was seen with the reduced expression of intracellular lipid and cholesterol transport proteins, *Plin4* and *Ncp1*. Like the CDI model, expression of *APOA4* was decreased in the hCE when exposed to toxins indicating that inhibition of cholesterol transport by *C. difficile* is dependent on toxins. Modification of cholesterol transport is seen in various infections to enhance pathogenesis. *Cryptosporidium parvum*, an intestinal parasite infecting ileal cells, reduces apolipoprotein expression and protein content in these cells to reduce cholesterol transport during infection^37^. Human cytomegalovirus (HCMV) modifies cholesterol transport by enhancing cholesterol efflux rather than inhibition. HCMV mobilizes lipid rafts via manipulation of actin filaments to facilitate cholesterol binding to APOA1 and increasing cholesterol efflux^38^.

Despite no significant changes in expression in FXR BA regulatory genes in the CDI mouse model, we did observe significant changes when exposed to toxins TcdA and TcdB in the two different colonic epithelial cell types. Caco-2 and hCE expression data show increased expression of BA regulatory genes *NR0B2* and *FGF19*. These proteins are important not only in BA regulation but also have a role in lipid and cholesterol metabolism. SHP (*NR0B2*) is an orphan response regulator that binds to transcriptional regulatory proteins altering their ability to initiate transcription. FGF19 enhances SHP activity through phosphorylation of SHP at T58. This partnership between SHP and FGF19 is able to modify expression of key cholesterol and BA transport genes^39,40^. The apical sodium bile salt transporter gene *SLC10A2* (ASBT) is down regulated through SHP repression of the LRH-1 transcriptional regulator^41,42^. The increase in *NR0B2* and *FGF19* expression and the significant reduction in *SLC10A2* expression in hCE cells suggest that toxins initiate the SHP/FGF19 partnership to reduce influx of BAs into the cells by repressing *SLC10A2* expression and may explain increased concentrations of TCA in the intestinal lumen in early infection. The SHP/FGF19 partnership is also important in downregulating cholesterol transport into cells by repressing SREBF2, a major cholesterol metabolic transcriptional regulator and therefore inhibiting transcription of *Ncp1l1*, a major intestinal cholesterol transporter^39^. SHP is also a repressor of MTP, an APOB chaperone, and APOA4 integral to chylomicron assembly^43^.

Interestingly, *FABP6* gene expression also was significantly reduced in hCE when exposed to toxins. FABP6 is an intracellular BA transporter and is highly expressed during FXR activation. The reduced expression supports the reduced BA transport during toxin exposure. However, this reduction is independent of FXR activation as OCA exposure to hCE cells enhances *FABP6* expression. This reduction when exposed to toxins is mediated by an unknown mechanism. Similarly, FXR activation is known to increase *NR0B2* expression^44^, however the increased expression of *NR0B2* in this study is independent of FXR activation as OCA exposure does not increase expression of *NR0B2* in the hCE cells. These results suggest that BA transport can be modified by *C. difficile* in a toxin dependent manner.

Repression of BA transport may be advantageous for *C. difficile* by effectively increasing the concentrations of BA in the lumen to enhance spore germination or growth. BA transport has been observed in the presence of other toxins and pathogens. The fungal mycotoxin deoxynivalenol alters BA transport by inhibiting *SLC10A2 FABP6* and *SLC51A* (OSTα)^45^. Similarly, infection with *Salmonella Typhimurium* in the porcine ileum reduces expression of *SLC10A2* and *FABP6*^46^. The increase in *FGF19* also suggests that *C. difficile* toxins not only can inhibit BA transport but may also have the potential to disrupt BA synthesis in the liver since FGF19 is a signaling protein that is transported to the liver to initiate reduction of BA synthesis.

*C. difficile* toxins are able to promote epithelial disruption by inhibiting the Rac and Rho GTPase signaling pathways that maintain cytoskeletal structure through glucosylating the GTPase via the GTD domain of the toxins^6-8^. This disruption also promotes altered expression of FXR regulated genes (Fig. 2A. Supp. Fig. 2) suggesting that toxins are facilitating a complex response in epithelial cells via GTPase disruption to promote disease and the success of *C. difficile*.

Three different platforms were utilized to interrogate the effects *C. difficile* toxins have on BA regulatory pathways. Gene expression in the CDI mouse model did not result in changes in FXR regulatory genes as was observed in the tissue culture experiments. This may be due to the high levels of the BA TβMCA found in the mouse inhibiting FXR activation. The BA pool was not measured in this study and may have an unknown impact. The presence of other microbes as well as the host immune response complicate our understanding. Tissue culture studies are advantageous as they reduce complicated interactions of an *in vivo* system. However, Caco-2 cell lines are derived from colonic cancer cells which inherently have altered gene expression and metabolism where results need to be interpreted cautiously. Interestingly, the hCE expression data obtained are more like the expression observed in the CDI mouse model then expression observed in Caco-2 cells, supporting previous work identifying hCE as a better mimic of cells *in vivo*^47^. Again, caution is needed as stem cells obtained from another individual may respond differently.

## Figure legends

**Supplemental Figure 1: Individual volcano plots of genes that changed in expression during *C. difficile* infection compared to antibiotic treated mice**. Volcano plots of genes differentially expressed in CDI mice compared to Cef mice. KEGG pathways are labeled in A) IL-17 signaling, B) Glycolosis/Gluconeogenesis, C) Cholesterol metabolism, and D) PPAR signaling pathway. E) Gene set enrichment analysis (GSEA) identifying gene clusters that changed the most in CDI compared to Cef mice.

**Supplemental Figure 2: TcdB alters FXR regulatory gene expression by disrupting GTPase signaling**. Caco-2 wells were treated with 100 pM of either TcdB, or 100 pM of TcdB-GTP^mut^ and incubated for 24 hr. NT, no treatment control only contained media with no other additions. Asterisks denote statistical significance determined by Mann-Whitney t-test compared to the No treatment control, \*\**p* < 0.01.

**Supplemental Table 2:** Probes used in NanoString analysis

**Supplemental Table 1:** Additional genes used in NanoString analysis.

| Gene Name | Protein Name | Significant Differential Expression in cecum (Log<sub>2</sub> fold change) |
| --- | --- | --- |
| **Nuclear receptors** |  |  |
| *Nr1i1* | VDR | Yes (-1.03) |
| *Nr1i2* | PXR | No |
| *Nr1i3* | CAR | Yes (-1.94) |
| *Rxra* | RXRA | Yes (-0.697) |
| *Gpbar1* | TGF5 | Not detected |
| *Nr1h2* | LXRb | Yes (-0.528) |
| *Nr1h3* | LXRa | Yes (-0.672) |
| *Nr1h4* | FXR | No |
| *Nr0b2* | SHP | Not detected |
| **Intestinal BA transport** |  |  |
| *Slc10a2* | ASBT | No |
| *Slc51a* | OSTa | Yes (-1.69) |
| *Slc51b* | OSTb | No |
| *Fabp6* | IBABP, FABP6 | No |
| *Abcc2* | ABCC2 | Not detected |
| *Slc9a3* | NHE3 | No |
| **Enterohepatic BA signaling** |  |  |
| *Fgf15* | FGF15 | Not detected |
| **Liver BA transport** |  |  |
| *Baat* | BAAT | Not detected |
| *Slc10a1* | NTCP | Not detected |
| **Liver BA transmembrane receptors** |  |  |
| *Klb* | b-Klotho | Not detected |
| *Fgfr4* | FGFR4 | No |
| *Vldlr* | VLDLR | No |
| *ldlr* | LDLR | No |
| **BA synthesis** |  |  |
| *Cyp27a1* | CYP27a1 | Yes (-1.99) |
| *Cyp7a1* | CYP7a1 | Not detected |
| *Cyp7b1* | CYP7b1 | No |
| *Cyp8b1* | CYP8b1 | No |

## References

1 Feuerstadt, P., Theriault, N. & Tillotson, G. The burden of CDI in the United States: a multifactorial challenge. BMC Infect Dis 23, 132 (2023). 10.1186/s12879-023-08096-0

2 Smits, W. K., Lyras, D., Lacy, D. B., Wilcox, M. H. & Kuijper, E. J. Clostridium difficile infection. Nat Rev Dis Primers 2, 16020 (2016). 10.1038/nrdp.2016.20

3 Guh, A. Y. et al. Trends in U.S. Burden of Clostridioides difficile Infection and Outcomes. N Engl J Med 382, 1320–1330 (2020). 10.1056/NEJMoa1910215

4 Slimings, C. & Riley, T. V. Antibiotics and hospital-acquired Clostridium difficile infection: update of systematic review and meta-analysis. J Antimicrob Chemother 69, 881–891 (2014). 10.1093/jac/dkt477

5 Mcdonald, L. C. et al. Clinical Practice Guidelines for Clostridium difficile Infection in Adults and Children: 2017 Update by the Infectious Diseases Society of America (IDSA) and Society for Healthcare Epidemiology of America (SHEA). Clinical Infectious Diseases 66, e1–e48 (2018). 10.1093/cid/cix1085

6 Aktories, K., Schwan, C. & Jank, T. Clostridium difficile Toxin Biology. Annual Review of Microbiology 71, 281–307 (2017). 10.1146/annurev-micro-090816-093458

7 Chumbler, N. M., Farrow, M. A., Lapierre, L. A., Franklin, J. L. & Lacy, D. B. Clostridium difficile Toxins TcdA and TcdB Cause Colonic Tissue Damage by Distinct Mechanisms. Infect Immun 84, 2871–2877 (2016). 10.1128/IAI.00583-16

8 Cowardin, C. A. et al. Glucosylation Drives the Innate Inflammatory Response to Clostridium difficile Toxin A. Infect Immun 84, 2317–2323 (2016). 10.1128/IAI.00327-16

9 Abt, M. C., McKenney, P. T. & Pamer, E. G. Clostridium difficile colitis: pathogenesis and host defence. Nat Rev Microbiol 14, 609–620 (2016). 10.1038/nrmicro.2016.108

10 Sorg, J. A. & Sonenshein, A. L. Bile salts and glycine as cogerminants for Clostridium difficile spores. J Bacteriol 190, 2505–2512 (2008). 10.1128/JB.01765-07

11 Theriot, C. M., Bowman, A. A. & Young, V. B. Antibiotic-Induced Alterations of the Gut Microbiota Alter Secondary Bile Acid Production and Allow for Clostridium difficile Spore Germination and Outgrowth in the Large Intestine. mSphere 1 (2016). 10.1128/mSphere.00045-15

12 Thanissery, R., Winston, J. A. & Theriot, C. M. Inhibition of spore germination, growth, and toxin activity of clinically relevant C. difficile strains by gut microbiota derived secondary bile acids. Anaerobe 45, 86–100 (2017). 10.1016/j.anaerobe.2017.03.004

13 Seekatz, A. M. et al. Fecal Microbiota Transplantation Eliminates Clostridium difficile in a Murine Model of Relapsing Disease. Infect Immun 83, 3838–3846 (2015). 10.1128/IAI.00459-15

14 McMillan, A. S. et al. Metagenomic, metabolomic, and lipidomic shifts associated with fecal microbiota transplantation for recurrent Clostridioides difficile infection. bioRxiv (2024). 10.1101/2024.02.07.579219

15 Wang, Y. D., Chen, W. D., Moore, D. D. & Huang, W. FXR: a metabolic regulator and cell protector. Cell Res 18, 1087–1095 (2008). 10.1038/cr.2008.289

16 Anderson, K. M. & Gayer, C. P. The Pathophysiology of Farnesoid X Receptor (FXR) in the GI Tract: Inflammation, Barrier Function and Innate Immunity. Cells 10 (2021). 10.3390/cells10113206

17 Fiorucci, S., Biagioli, M., Zampella, A. & Distrutti, E. Bile Acids Activated Receptors Regulate Innate Immunity. Front Immunol 9, 1853 (2018). 10.3389/fimmu.2018.01853

18 Gadaleta, R. M. et al. Activation of bile salt nuclear receptor FXR is repressed by pro-inflammatory cytokines activating NF-kappaB signaling in the intestine. Biochim Biophys Acta 1812, 851–858 (2011). 10.1016/j.bbadis.2011.04.005

19 Winston, J. A. et al. Ursodeoxycholic Acid (UDCA) Mitigates the Host Inflammatory Response during Clostridioides difficile Infection by Altering Gut Bile Acids. Infect Immun 88 (2020). 10.1128/IAI.00045-20

20 Makishima, M. et al. Identification of a Nuclear Receptor for Bile Acids. Science 284 (1999-5-21). 10.1126/science.284.5418.1362

21 Wexler, A. G. et al. Clostridioides difficile infection induces a rapid influx of bile acids into the gut during colonization of the host. Cell Rep 36, 109683 (2021). 10.1016/j.celrep.2021.109683

22 Winston, J. A., Thanissery, R., Montgomery, S. A. & Theriot, C. M. Cefoperazone-treated Mouse Model of Clinically-relevant Clostridium difficile Strain R20291. J Vis Exp (2016). 10.3791/54850

23 Theriot, C. M. et al. Cefoperazone-treated mice as an experimental platform to assess differential virulence of Clostridium difficile strains. Gut Microbes 2, 326–334 (2011). 10.4161/gmic.19142

24 Pike, C. M. et al. Characterization and optimization of variability in a human colonic epithelium culture model. ALTEX - Alternatives to animal experimentation 41, 425–438 (2024). 10.14573/altex.2309221

25 Wang, Y. et al. Self-renewing Monolayer of Primary Colonic or Rectal Epithelial Cells. Cell Mol Gastroenterol Hepatol 4, 165–182 e167 (2017). 10.1016/j.jcmgh.2017.02.011

26 Yang, G. et al. Expression of recombinant Clostridium difficile toxin A and B in Bacillus megaterium. BMC Microbiology 8, 192 (2008). 10.1186/1471-2180-8-192

27 Pike, C. M., Tam, J., Melnyk, R. A. & Theriot, C. M. Tauroursodeoxycholic Acid Inhibits Clostridioides difficile Toxin-Induced Apoptosis. Infection and Immunity 90 (2022-7-7). 10.1128/iai.00153-22

28 Sayin, S. I. et al. Gut microbiota regulates bile acid metabolism by reducing the levels of tauro-beta-muricholic acid, a naturally occurring FXR antagonist. Cell Metab 17, 225–235 (2013). 10.1016/j.cmet.2013.01.003

29 Ohmachi, T. et al. Fatty acid binding protein 6 is overexpressed in colorectal cancer. Clin Cancer Res 12, 5090–5095 (2006). 10.1158/1078-0432.CCR-05-2045

30 Gottardi, D. A. et al. The Bile Acid Nuclear Receptor FXR and the Bile Acid Binding Protein IBABP Are Differently Expressed in Colon Cancer. Digestive Diseases and Sciences 49, 982–989 (2004). 10.1023/B:DDAS.0000034558.78747.98

31 McMillan, A. S. & Theriot, C. M. Bile acids impact the microbiota, host, and C. difficile dynamics providing insight into mechanisms of efficacy of FMTs and microbiota-focused therapeutics. Gut Microbes 16 (2024-12-31). 10.1080/19490976.2024.2393766

32 Hussain, M. M. et al. Chylomicron assembly and catabolism: role of apolipoproteins and receptors. Biochimica et Biophysica Acta (BBA) - Lipids and Lipid Metabolism 1300 (1996/05/20). 10.1016/0005-2760(96)00041-0

33 Kohan, A. B., Wang, F., Lo, C.-M., Liu, M. & Tso, P. ApoA-IV: current and emerging roles in intestinal lipid metabolism, glucose homeostasis, and satiety. American Journal of Physiology-Gastrointestinal and Liver Physiology 308 (2015 Mar 15). 10.1152/ajpgi.00098.2014

34 Ko, C.-W., Qu, J., Black, D. D. & Tso, P. Regulation of intestinal lipid metabolism: current concepts and relevance to disease. Nature Reviews Gastroenterology & Hepatology 17, 169–183 (2020). 10.1038/s41575-019-0250-7

35 Colin, S. et al. Activation of intestinal peroxisome proliferator-activated receptor-increases high-density lipoprotein production. European Heart Journal 34, 2566–2574 (2013). 10.1093/eurheartj/ehs227

36 Guardiola, M. et al. APOA5 gene expression in the human intestinal tissue and its response to in vitro exposure to fatty acid and fibrate. Nutrition, Metabolism and Cardiovascular Diseases 22 (2012/09/01). 10.1016/j.numecd.2010.12.00337

37 Wang, S. et al. (2024).

38 Low, H. et al. Cytomegalovirus Restructures Lipid Rafts via a US28/CDC42-Mediated Pathway, Enhancing Cholesterol Efflux from Host Cells. Cell Rep 16, 186–200 (2016). 10.1016/j.celrep.2016.05.070

39 Kim, Y. C. et al. Small Heterodimer Partner and Fibroblast Growth Factor 19 Inhibit Expression of NPC1L1 in Mouse Intestine and Cholesterol Absorption. Gastroenterology 156, 1052–1065 (2019). 10.1053/j.gastro.2018.11.061

40 Kim, Y. C. et al. Transgenic mice lacking FGF15/19-SHP phosphorylation display altered bile acids and gut bacteria, promoting nonalcoholic fatty liver disease. J Biol Chem 299, 104946 (2023). 10.1016/j.jbc.2023.104946

41 Chen, F. et al. Liver receptor homologue-1 mediates species- and cell line-specific bile acid-dependent negative feedback regulation of the apical sodium-dependent bile acid transporter. J Biol Chem 278, 19909–19916 (2003). 10.1074/jbc.M207903200

42 Nguyen, J. T., Riessen, R., Zhang, T., Kieffer, C. & Anakk, S. Deletion of Intestinal SHP Impairs Short-term Response to Cholic Acid Challenge in Male Mice. Endocrinology 162 (2021). 10.1210/endocr/bqab063

43 Pan, X. & Hussain, M. M. Bmal1 regulates production of larger lipoproteins by modulating cAMP-responsive element-binding protein H and apolipoprotein AIV. Hepatology 76, 78–93 (2022). 10.1002/hep.32196

44 Vavassori, P., Mencarelli, A., Renga, B., Distrutti, E. & Fiorucci, S. The Bile Acid Receptor FXR Is a Modulator of Intestinal Innate Immunity. The Journal of Immunology 183, 6251–6261 (2009). 10.4049/jimmunol.0803978

45 Wang, J. et al. Exposure to the mycotoxin deoxynivalenol reduces the transport of conjugated bile acids by intestinal Caco-2 cells. Arch Toxicol 96, 1473–1482 (2022). 10.1007/s00204-022-03256-8

46 Uribe, J. H. et al. Transcriptional analysis of porcine intestinal mucosa infected with Salmonella Typhimurium revealed a massive inflammatory response and disruption of bile acid absorption in ileum. Veterinary Research 47 (2016). 10.1186/s13567-015-0286-9

47 Ok, M. T. et al. A leaky human colon model reveals uncoupled apical/basal cytotoxicity in early Clostridioides difficile toxin exposure. Am J Physiol Gastrointest Liver Physiol 324, G262–G280 (2023). 10.1152/ajpgi.00251.2022

